# Microbial metagenome of urinary tract infection

**DOI:** 10.1101/134320

**Authors:** Ahmed Moustafa, Harinder Singh, Weizhong Li, Kelvin J. Moncera, Manolito G. Torralba, Yanbao Yu, Oriol Manuel, William Biggs, J. Craig Venter, Karen E. Nelson, Rembert Pieper, Amalio Telenti

**Author notes:** Corresponding Authors: Amalio Telenti, Rembert Pieper. Emails authors: Ahmed Moustafa < >, Singh, Harinder < >, Weizhong Li, Torralba, Manolito, Yu, Yanbao < >, Manuel Josep Oriol < >, William Biggs < >, Craig Venter, Karen Nelson < >, Pieper, Rembert < >.

## Abstract

Urine culture and microscopy techniques are used to profile the bacterial species present in urinary tract infections. To gain insight into the urinary flora in infection and health, we analyzed clinical laboratory features and the microbial metagenome of 121 clean-catch urine samples. 16S rDNA gene signatures were successfully obtained for 116 participants, while whole genome shotgun sequencing data was successfully generated for samples from 49 participants. Analysis of these datasets supports the definition of the patterns of infection and colonization/contamination. Although 16S rDNA sequencing was more sensitive, whole genome shotgun sequencing allowed for a more comprehensive and unbiased representation of the microbial flora, including eukarya and viral pathogens, and of bacterial virulence factors. Urine samples positive by whole genome shotgun sequencing contained a plethora of bacterial (median 41 genera/sample), eukarya (median 2 species/sample) and viral sequences (median 3 viruses/sample). Genomic analyses revealed cases of infection with potential pathogens (e.g., *Alloscardovia sp, Actinotignum sp*, *Ureaplasma sp*) that are often missed during routine urine culture due to species specific growth requirements. We also observed gender differences in the microbial metagenome. While conventional microbiological methods are inadequate to identify a large diversity of microbial species that are present in urine, genomic approaches appear to comprehensively and quantitatively describe the urinary microbiome.

## Introduction

Urinary tract infections (UTIs) occur in a high proportion of the population and are a significant health economic burden [1]. The criteria for diagnosis includes multiple clinical parameters and laboratory tests [2], and the clinical suspicion of a UTI frequently triggers the prescription of broad spectrum antibiotics, with or without confirmation of the infecting organisms. The most common organism in uncomplicated UTIs is *Escherichia coli* followed by a number of gram-positive cocci and other Enterobacteriaceae [3]. Other organisms, including difficult-to-culture prokaryotes, eukaryotes such as *Candida albicans* and viruses, are involved in UTIs or other manifestations of genitourinary tract infection such as urethritis and sexually transmitted diseases. Because the care of UTI is streamlined, it is only after treatment failure that molecular tests and additional non-molecular investigations are launched.

Conventional microbiological methods are inadequate to fully determine the diversity of bacteria that are present in urine [4]. Next generation sequencing techniques create the possibility of investigating the microbial metagenome associated with infection and inflammation of the urinary tract. Metaproteomic methods have enabled a deeper characterization of the inflammatory response towards uropathogens in cases of UTI and asymptomatic bacteriuria [5, 6]. Several studies have characterized the urinary microbiome by 16S rRNA sequencing to study adults with a cumulative number of 253 subjects investigated across these studies (UTI, n=50; other urinary manifestations, n=154; sexually transmitted diseases, n=20; renal transplant samples, n=60) [4, 7–19]. However, a complete view of the microbiome, including eukarya and viruses, as well as an unbiased characterization of abundance and the identification of virulence factors as presented in this study can only be achieved by comprehensive metagenome analyses using whole genome shotgun sequencing (WGS). A preliminary study of the urine metagenome was recently published by Hasman and colleagues [20] and hinted at the increased microbial diversity observed using unbiased sequencing. Ion torrent was the technology used in the Hasman et al. study.

The aims of this study were to discover new microbial and viral components in clinical urine specimens using metagenomics sequencing combined with 16S rDNA and metaproteomic analyses, and to examine the question of whether the microbial compositions of urine specimens justifies the description of a urinary microbial metagenome. The metagenomics component allowed for the exploration of organisms and their abundances from all microbial kingdoms and allowed us to investigate the distribution of known virulence genes across the various study groups. Overall, the study reveals patterns of peri-urethral colonization and vaginal contamination of urine samples and of different profiles of what can be considered active infection groups. The study also contributes to the identification of difficult-to-culture and potential novel pathogens and addresses the presence of various human viruses and eukarya that are important in genitourinary medicine.

## Results

### Clinical and laboratory data representation

To support an unbiased assessment of the clinical nature of the specimens, we approached the urine sample laboratory and microbiology data using dimensionality reduction, and K-mer clustering analysis. The PCA representation of the clinical laboratory data is presented in Fig 1. The PCA analysis showed that the first two components (PC1, PC2) explained 65% of the variance in the clinical laboratory dataset. PC1 was driven by the vaginal contamination score (VCO), PC2 was contributed primarily by neutrophil activation and degranulation score (NAD), and secondarily by the erythrocyte and vascular injury score (ERY) and the presence of red blood cells (RBC) and leukocytes (WBC) (Fig 1B). The partitioning around medoids clustering resulted in three Clusters, with 9 individuals in Cluster #1, 63 individuals in Cluster #2, and 49 individuals in Cluster #3 (Fig 1C). From these data, we established a preliminary definition of Cluster #1 as likely representing urine from non-infected individuals, while Clusters #2 and #3 are consistent with separate manifestations of infectious and inflammatory processes of the urinary tract. The performance of 16S rDNA and whole genome shotgun sequencing across clinical laboratory clusters is presented in Table 1.

**Table 1.** Microbiome Sequencing performance rate.

|  | Cluster 1 |  | Cluster 2 |  | Cluster 3 |  | Total |  |
| --- | --- | --- | --- | --- | --- | --- | --- | --- |
| Microbiome | Count | Percent | Count | Percent | Count | Percent | Count | Percent |
| **WGS+16S** | 5 | 56% | 35 | 56% | 7 | 14% | 47 | 39% |
| **WGS** | 0 | 0% | 1 | 2% | 1 | 2% | 2 | 2% |
| **16S** | 4 | 44% | 27 | 43% | 38 | 78% | 69 | 57% |
| **None** | 0 | 0% | 0 | 0% | 3 | 6% | 3 | 2% |
| **Total** | 9 |  | 63 |  | 49 |  | 121 |  |

WGS: whole genome shotgun sequencing, 16S: 16S rDNA sequencing

**Figure 1.**
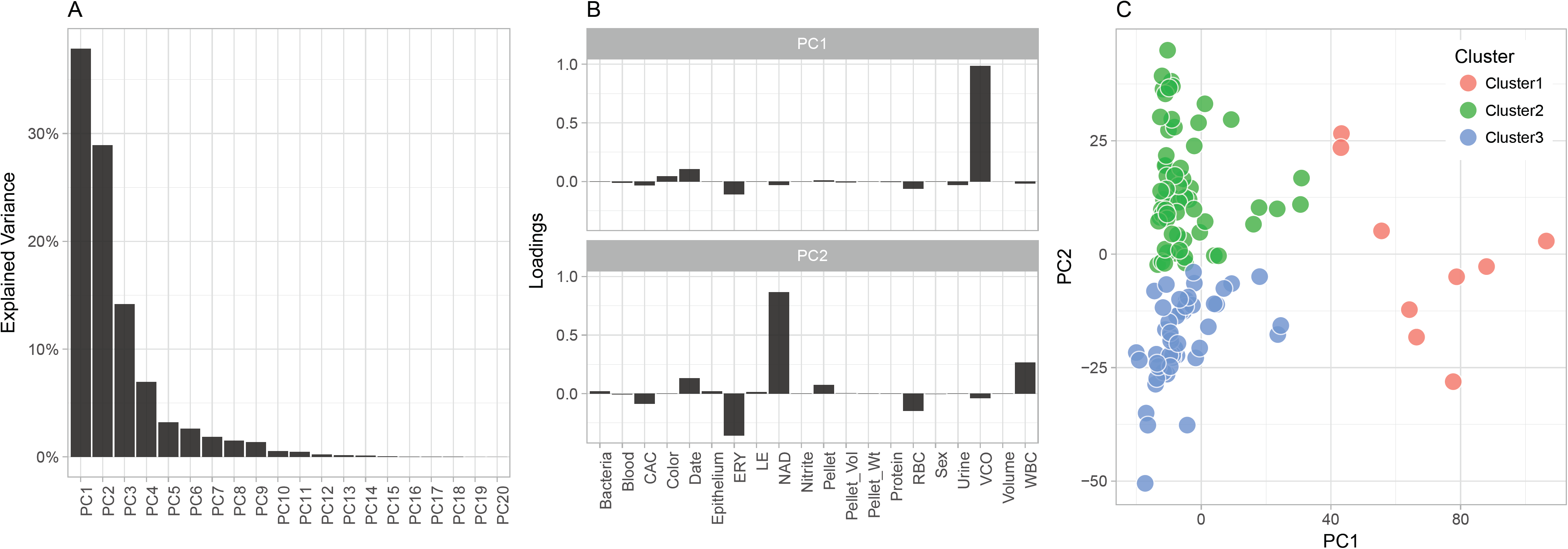
Definition of clinical and laboratory groups. The study used an unbiased approach to the classification of specimens using 20 parameters from the laboratory analysis of urine. (A) Explained variance from PCA; the first two PCs were retained for downstream clustering analyses. (B) Contributing factors (loadings) to the first two PCs. Note that the directionality of the loadings reflect enrichment independently of sign and direction. (C) Clustering of samples is based on the method of partitioning around medoids (pam).

**Figure 2.**
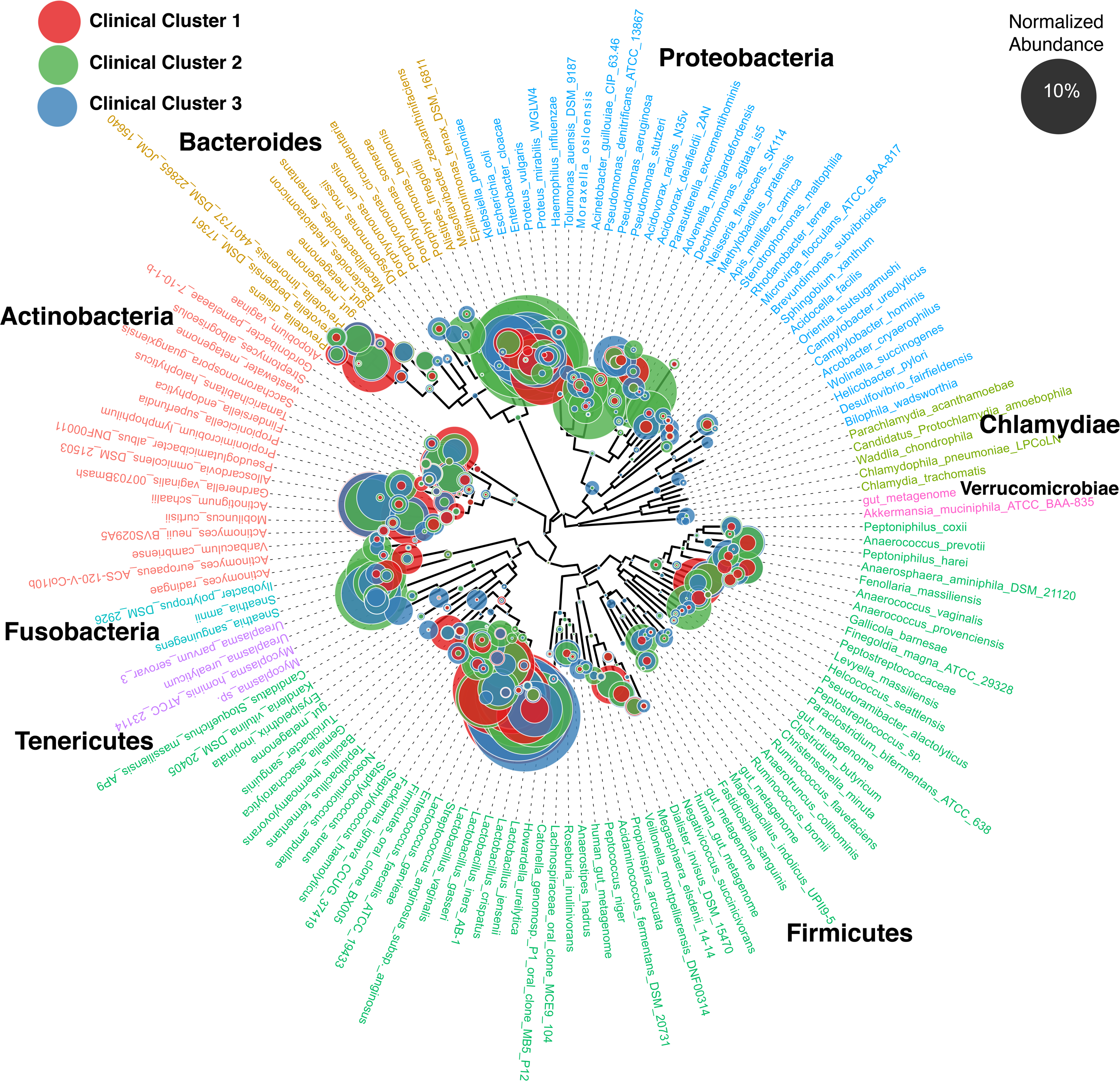
Normalized abundance of bacterial genera across the clinical groups using 16S rDNA. 116 samples were successfully analysed by 16S rDNA sequencing and grouped according to the clinical laboratory clusters. Proteobacteria were the predominant phylum in Cluster 2 - the cluster that represents infection, with prominent identification of *Citrobacter*, *Enterobacter*, *Escherichia*. Clusters 1 and 3 were more diverse in composition (S1 Fig).

### 16S rDNA sequencing

16S rDNA sequencing was successful for 116 (96%) samples (Table 1). The median (range) number of genera identified per individual was 38 (6-220). The median (range) number of genera varied across clinical Clusters: 51(16-106) for Cluster 1, 32 (7-172) for Cluster 2, and 60 (6-220) for Cluster 3. Analysis of the normalized abundance of the classified bacterial genera across the clinical groups confirmed that proteobacteria were the predominant phylum in cluster 2 - the Cluster that represents infection, with prominent identification of *Citrobacter sp.*, *Enterobacter sp.*, and *Escherichia sp*. Clusters 1 and 3 were more diverse in composition. Cluster 1 had greater abundance of *Actinotignum*, *Aerococcus*, *Atopobium*, *Facklamia*, *Gardnerella*, *Lactobacillus*, *Megasphaera*, *Oligella*, *Prevotella*, and *Streptococcus* species. Cluster 3 had greater abundance of *Acidovorax*, *Alloscardovia*, *Epilithonimonas*, *Lachnospira*, *Peptostreptococcus*, *Pseudomonas*, *Rhodanobacter*, *Riemerella*, *Sphingobium* and *Ureaplasma* (S1 Fig). Overall, 16S rDNA sequencing may prove too sensitive to define significant components of the urinary microbiome.

### Whole-genome shotgun sequencing

Shotgun metagenome analysis was successfully performed on 49 samples highlighting that, although samples with limited microbial content may amplify via 16S rDNA, insufficient reads or failed sequencing will occur if starting DNA material is limiting. However, metagenomics data will reflect quantitatively more accurate analyses compared to 16S rDNA data. In the samples that were successfully sequenced by WGS, the average composition of the reads per kingdom was 94.6% Bacteria, 0.05% Eukarya, 0.0027% Viruses, and 0.0001% Archaea (Fig 3). The archaeal component was discarded from subsequent analyses. We also observed a significant proportion, 4.9%, of unmapped, non-human sequence reads. The largest microbial content was observed in Cluster 1, the lowest in Cluster 3. This is an important observation that, together with other features (see below), could indicate that Cluster 3 represent urinary samples from individuals that received treatment.

**Figure 3.**
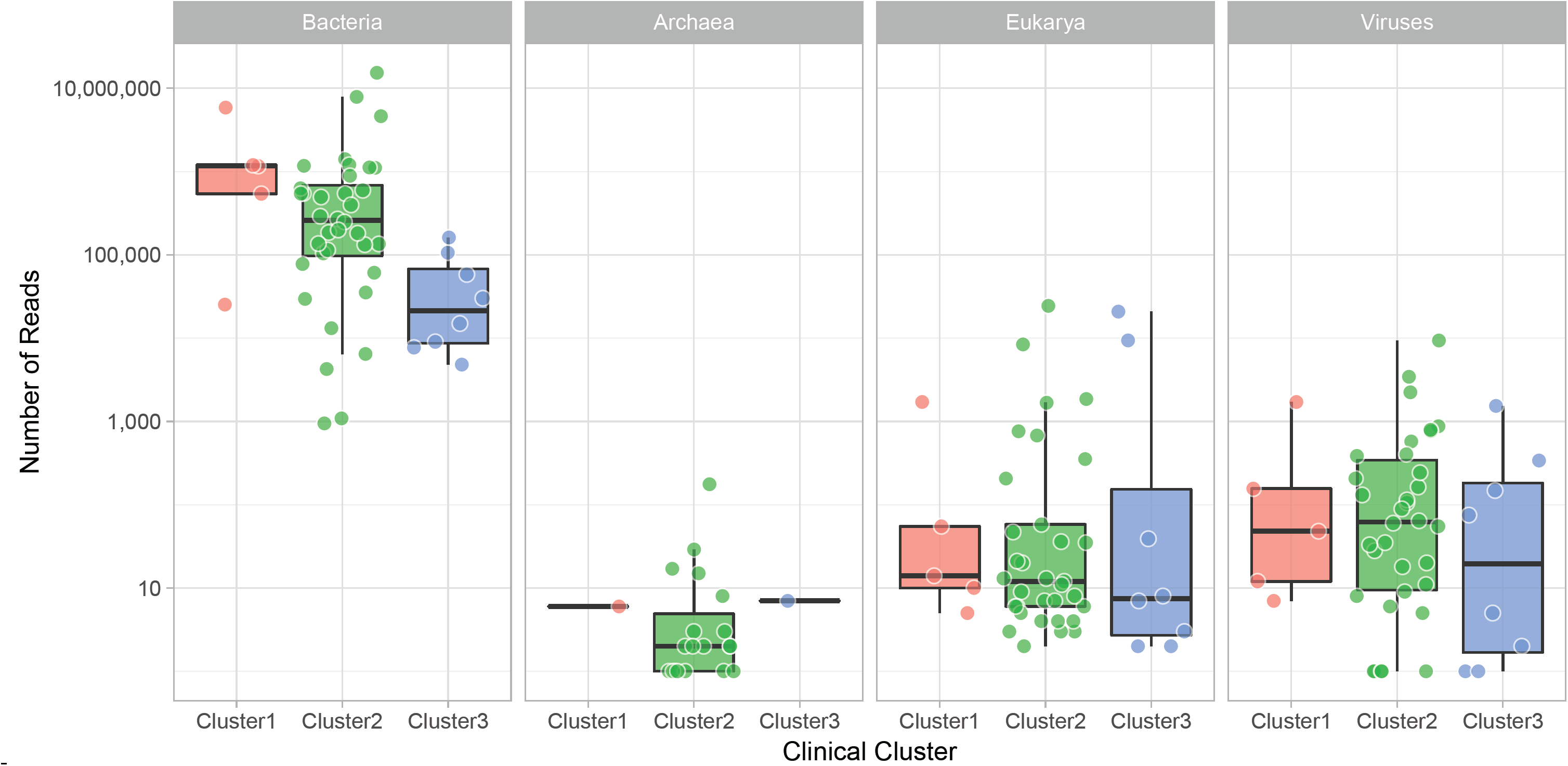
Whole genome shotgun sequencing mapped reads per sample. 49 samples were successfully sequenced and grouped according to the clinical laboratory clusters. Each point represents a sample. The thick line in the boxplot represents the median number of reads for the cluster.

### Bacteria

The median (range) number of bacterial species – genera - identified per individual was 41 (27-49). The median (range) number of species across clinical Clusters was 44 (29-48) for Cluster 1, 41 (28-49) for Cluster 2, and 38 (28-47) for Cluster 3. Fig 4 depicts the read counts for genera across clinical groups. This analysis indicates that proteobacteria are the predominant phylum in Cluster 2 - the cluster that represents infection, including classic uropathogens such as *Escherichia*, *Klebsiella*, *Pseudomonas*, *Enterobacter*, *Citrobacter,* as well as species with unclear or unknown role in infection, such as *Acidovorax*, *Rhodanobacter,* and *Oligella* (S2 Fig). Cluster 1 had greater abundance of *Actinomyces*, *Anaerococcus*, *Atopobium*, *Facklamia*, *Finegoldia*, *Gardnerella*, *Lactobacillus*, *Megasphaera*, *Peptoniphilus*, *Staphylococcus*, and *Streptococcus* (S2 Fig). Given the depletion in total number of reads, and the elevated diversity in species in Cluster 3, we could not identify any uniquely enriched genus.

**Figure 4.**
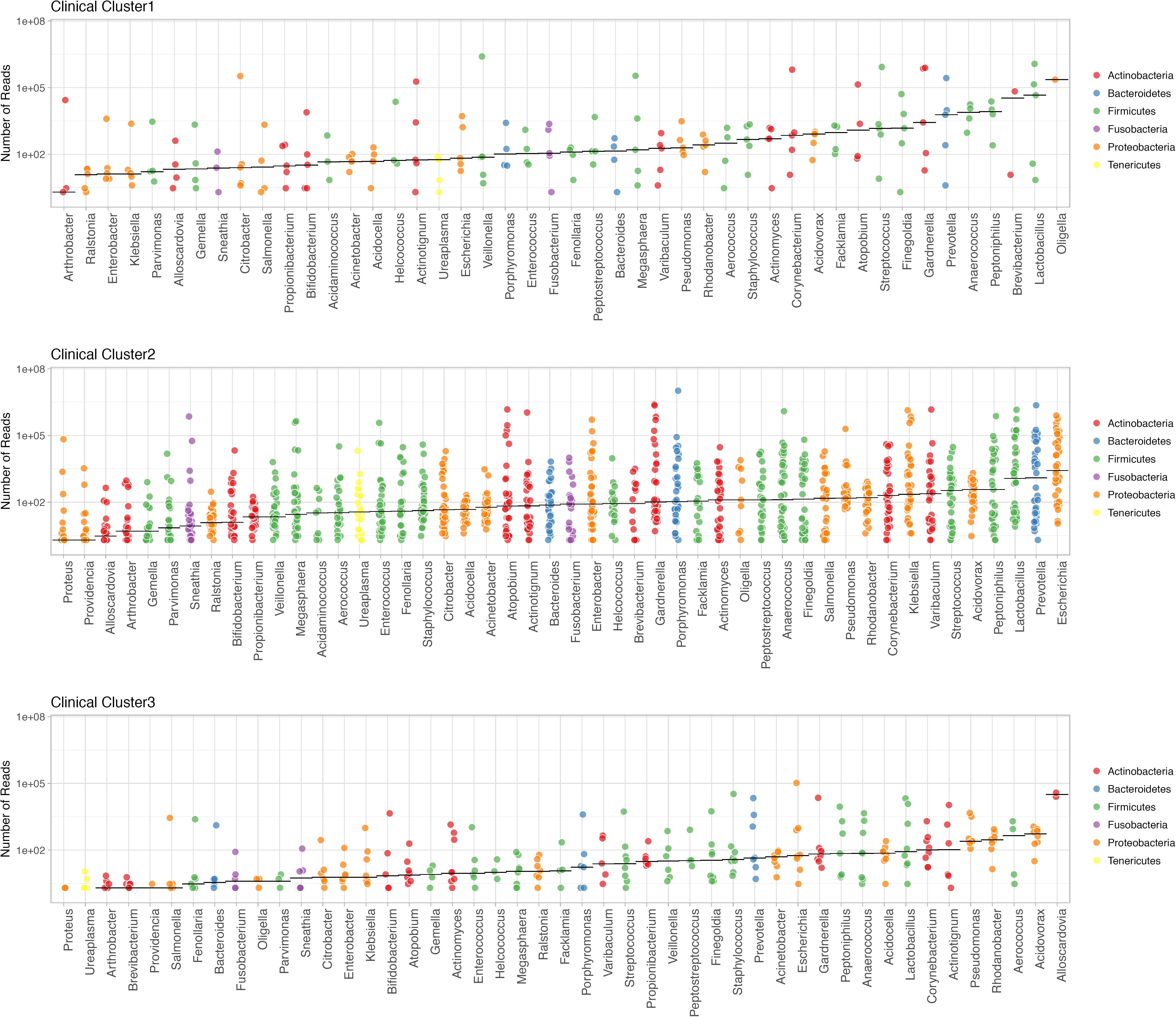
Ranking of bacterial genera by counts from whole genome shotgun sequencing across clinical laboratory clusters. Shown are bacteria observed with at least 1% of total reads in a sample. Analyses reflect results from 49 samples that were successfully sequenced and grouped according to the clinical laboratory clusters. Each point represents a genus in a sample. The horizontal represents the median number of reads for the genus.

We specifically chose to represent the metagenome data as read counts, a surrogate of absolute abundance, because the process does not involve amplification and thus, relative abundance may misrepresent actual content of microbiota. However, we compared the relative abundance as estimated by 16S rDNA with the absolute read number from metagenome sequencing to assess the degree of correlation. The correlation was high (R^2^=0.88), however, there were some discrepancies where relevant organisms appeared better identified by WGS than by 16S rDNA sequencing (eg. *Gardnerella*, S3 Fig). We also explored the nature of samples in Cluster 2 that were negative by WGS – despite of the expectation that samples in this group would be indicative of infection. For this, we inspected differences in 16S rDNA read counts for 27 samples in Cluster 2 that were negative in WGS compared with 35 samples in Cluster 2 that were positive in WGS. We did not identify differences in median 16S rDNA bacterial read counts across these two sets, nor a significant difference in pattern of bacterial abundance. Therefore, it remains unclear what the true nature of those Cluster 2 samples is: inflammatory reactions, traumatic (for example, passage of a kidney stone), low grade infection, or technical limits to WGS that limit sensitivity.

It was also important to assess the correspondence of WGS and routine culture used in the clinics. A total of 23 samples in Cluster 2 presented dominant flora (*post hoc* defined as ≥10^5^ reads). For those, we observed eight samples with consistent WGS and culture results, 1 with a discrepant growth, and 4 reported as mixed flora in culture (Table 2). There was no reported culture growth for four samples in Cluster 1, and only one sample in Cluster 3 despite the observation of dominant flora in sequencing. Two samples, one in Cluster 1(colonization/contamination) and Cluster 2 (infection) contained high number of reads of *Actinotignum* sp. This facultative anaerobic gram-positive rod (in particular, *A. schaali*) has been claimed to be part of the urinary microbiota of healthy individuals while also responsible for UTIs, particularly in elderly men and young children [21]. *A. schaalii* may be an underestimated cause of UTIs because of its fastidious growth on usual media and difficulties associated with its identification using phenotypic methods [21]. Use of matrix-assisted laser desorption/ionisation time-of-flight mass spectrometry (MALDI-TOF MS) supports the better identification of this organism [22].

**Table 2.**
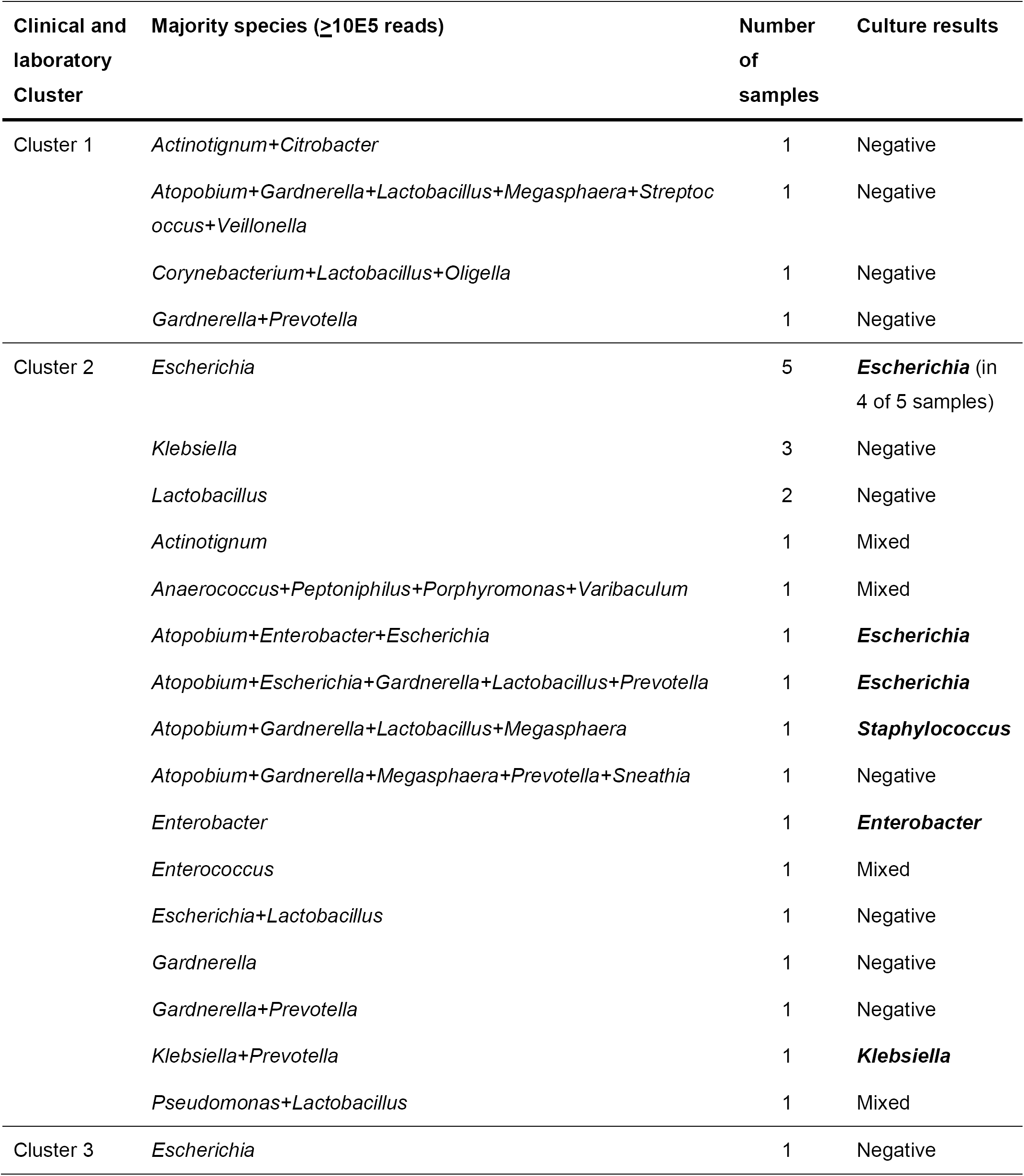
Relationship between whole genome shotgun sequencing and routine culture.

| Clinical and laboratory Cluster | Majority species ($\geq 10^5$ reads) | Number of samples | Culture results |
| --- | --- | --- | --- |
| Cluster 1 | *Actinotignum*+*Citrobacter* | 1 | Negative |
|  | *Atopobium*+*Gardnerella*+*Lactobacillus*+*Megasphaera*+*Streptococcus*+*Veillonella* | 1 | Negative |
|  | *Corynebacterium*+*Lactobacillus*+*Oligella* | 1 | Negative |
|  | *Gardnerella*+*Prevotella* | 1 | Negative |
| Cluster 2 | *Escherichia* | 5 | ***Escherichia*** (in 4 of 5 samples) |
|  | *Klebsiella* | 3 | Negative |
|  | *Lactobacillus* | 2 | Negative |
|  | *Actinotignum* | 1 | Mixed |
|  | *Anaerococcus*+*Peptoniphilus*+*Porphyromonas*+*Varibaculum* | 1 | Mixed |
|  | *Atopobium*+*Enterobacter*+*Escherichia* | 1 | ***Escherichia*** |
|  | *Atopobium*+*Escherichia*+*Gardnerella*+*Lactobacillus*+*Prevotella* | 1 | ***Escherichia*** |
|  | *Atopobium*+*Gardnerella*+*Lactobacillus*+*Megasphaera* | 1 | ***Staphylococcus*** |
|  | *Atopobium*+*Gardnerella*+*Megasphaera*+*Prevotella*+*Sneathia* | 1 | Negative |
|  | *Enterobacter* | 1 | ***Enterobacter*** |
|  | *Enterococcus* | 1 | Mixed |
|  | *Escherichia*+*Lactobacillus* | 1 | Negative |
|  | *Gardnerella* | 1 | Negative |
|  | *Gardnerella*+*Prevotella* | 1 | Negative |
|  | *Klebsiella*+*Prevotella* | 1 | ***Klebsiella*** |
|  | *Pseudomonas*+*Lactobacillus* | 1 | Mixed |
| Cluster 3 | *Escherichia* | 1 | Negative |

The metagenome approach permits the identification of virulence genes in the bacterial pool. Searching for virulence factors against VFDB [23], we observed enrichment of specific factors, in particular in Cluster 2 (Fig 5).

**Figure 5.**
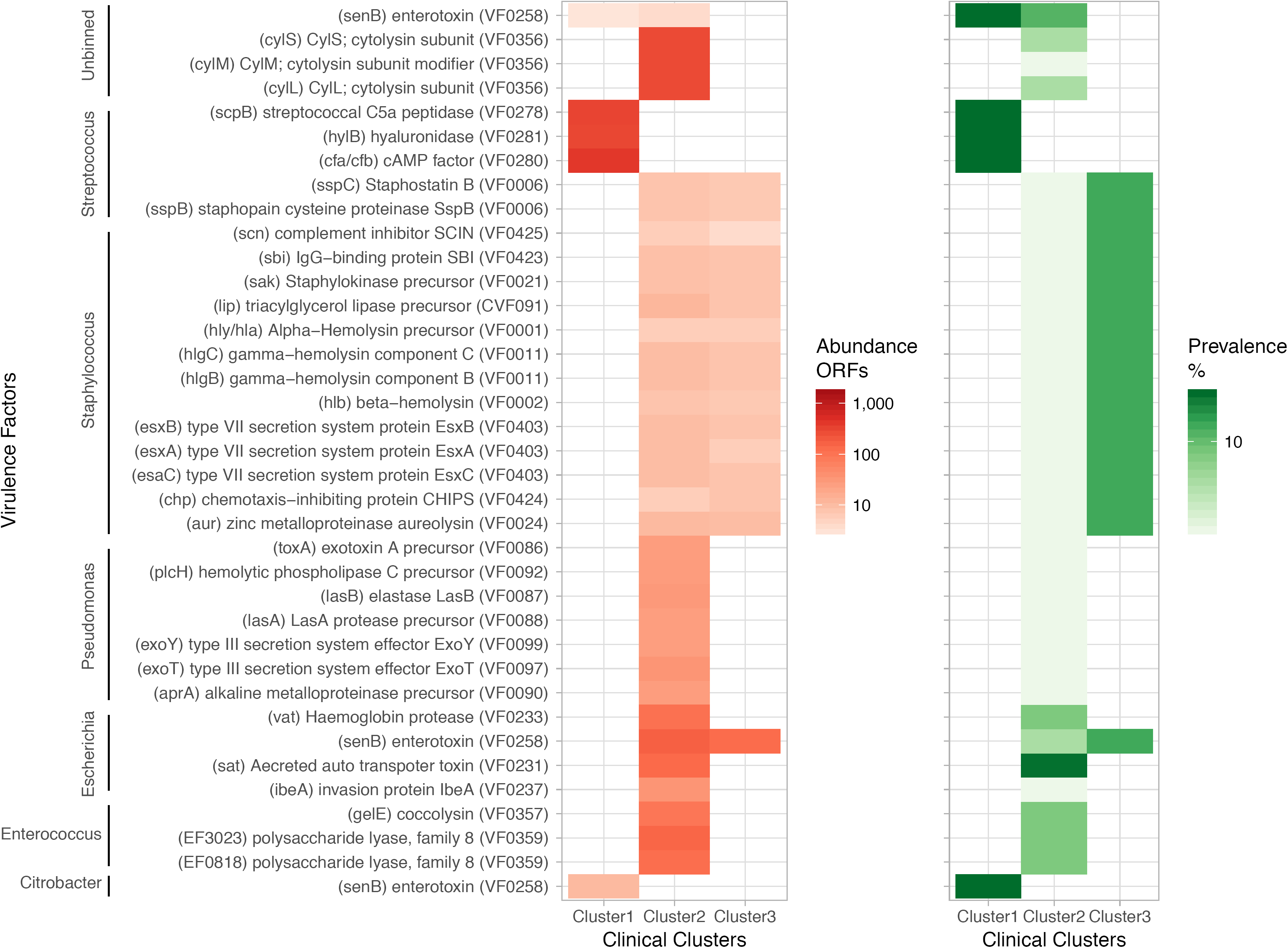
Virulence factors across clinical laboratory clusters. Whole-genome shotgun sequencing data was used to search for open reading frames (ORFs) compared against the database VFDB [23] to identify virulence factor genes with over 90% sequence identity. Listed are the factors identified in the dataset, group by taxonomic binning, with the VFDB accession number in parenthesis. The left panel shows enrichment in number of ORFs across clusters. The right panel shows prevalence of samples that contain organisms carrying the corresponding virulence factor in each cluster.

### Eukarya

The median (range) number of species identified per individual was 2 (1-8). The median (range) number of species was 2 (1-8) for cluster 1, 2 (1-6) for cluster 2, and 2 (1-3) for cluster 3. Nine species were identified (minimum 10 reads per sample): eight fungal species (*Candida albicans, C. grabrata, C. orthopsilosis* and *C. tropicalis, Clavispora lusitaniae, Lodderomyces elongisporus, Meyerozyma guilliermondii and Malassezia globosa*) and a metamonada (*Trichomonas vaginalis*). Fig 6 depicts the read counts for genera across clinical groups. Relatively elevated counts were observed for *C. glabrata* and *Clavispora lusitaniae* in four individuals from Clusters 2 and 3.

**Figure 6.**
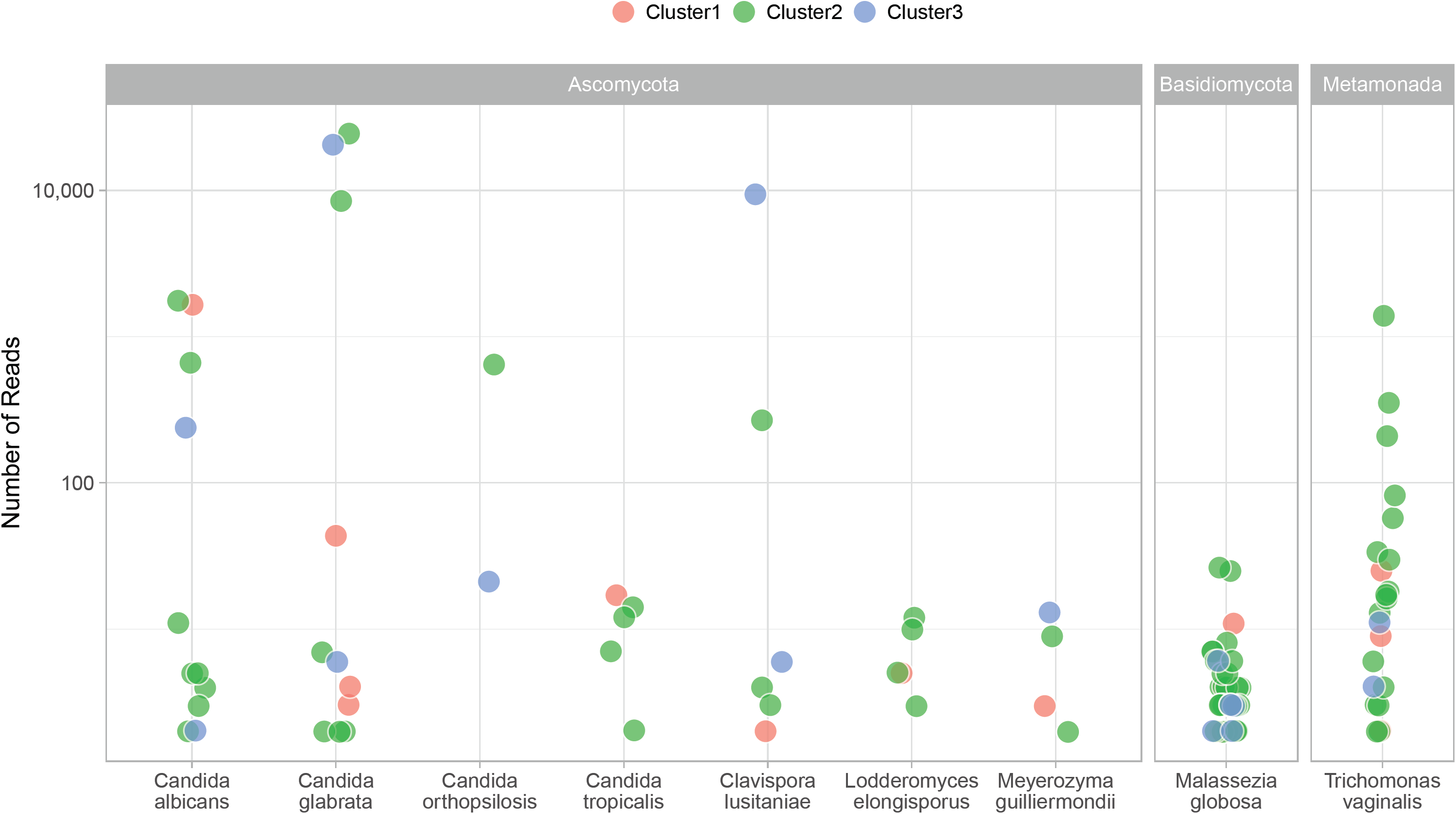
Eukarya read counts across clinical laboratory clusters. Shown are eukarya observed with at least 10 sequence reads in a sample. Analyses reflect results from 49 samples that were successfully sequenced and grouped according to the clinical laboratory clusters. Each point represents a species in a sample.

Candida species both colonize and cause invasive disease in the urinary tract [24]. The identification of the lipophilic fungi *Malassezia* is not unexpected as these fungi predominate in most skin sites in healthy adults [25].

*Trichomonas vaginalis* colonizes the genitourinary tract of men and women. Young women with urinary symptoms in the absence of documented UTI were found more likely to have *Trichomonas vaginalis* compared to those with a documented UTI [26]. Molecular amplification detects *Trichomonas vaginalis* in penile-meatal swabs and urine specimens of men [27]. The identification in the present study of sequence reads in 18 of 38 females (47%) and 4 of 11 (36%) males suggests the common presence of this organism in the genitourinary region.

### Viruses

The median (range) number of viruses identified per individual was 3 (1-9). The median (range) number of viruses was 3 (2-6) for cluster 1, 3 (1-9) for cluster 2, and 2 (1-7) for cluster 3 (Fig 7). We identified 13 phages that were generally dominant and associated with the cognate bacteria in the sample

**Figure 7.**
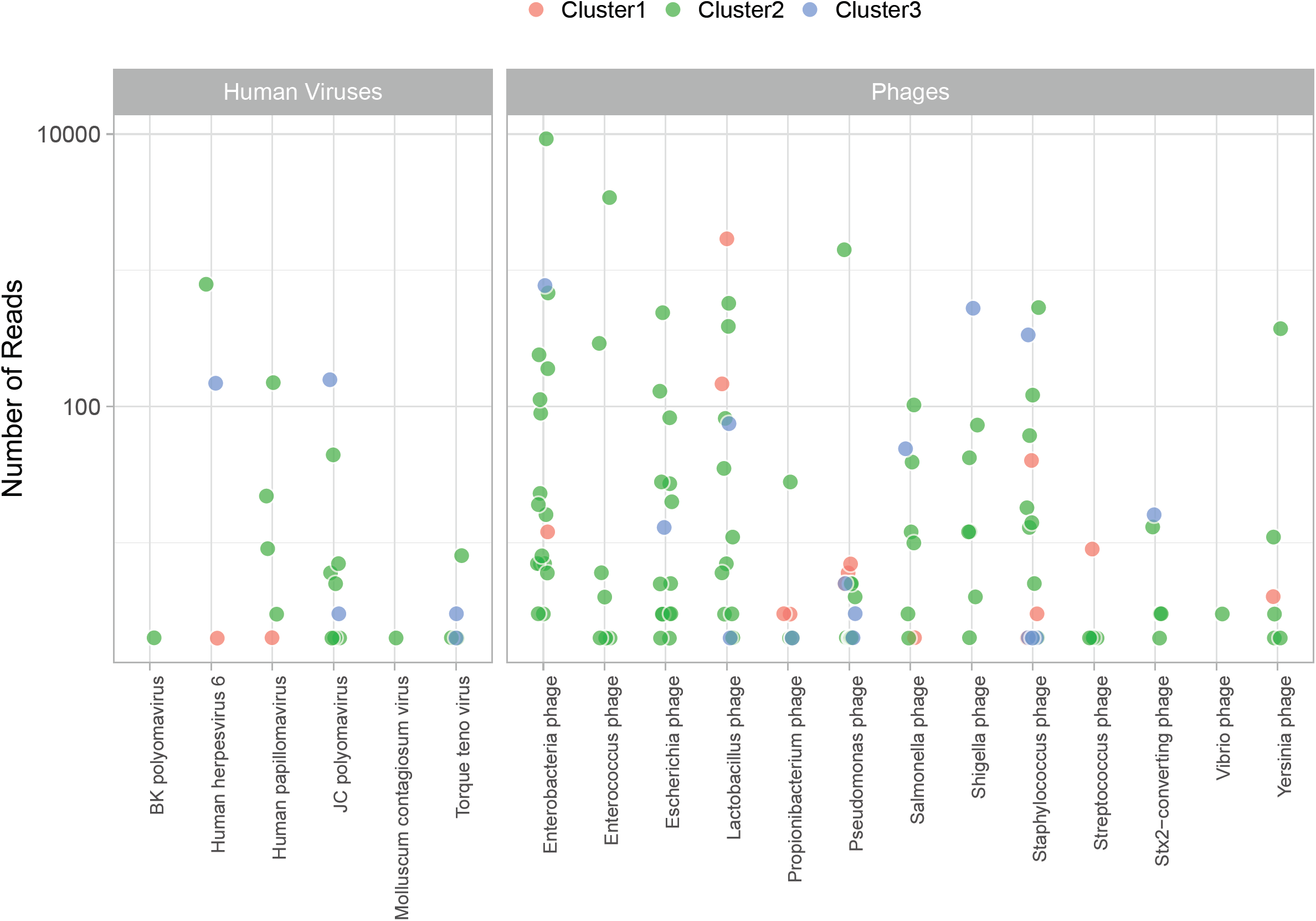
Viral read counts across clinical laboratory clusters. Shown are viruses observed with at least one sequence read in a sample. Analyses reflect results from 49 samples that were successfully sequenced and grouped according to the clinical laboratory clusters. Each point represents a virus in a sample.

We identified 6 human viruses consistent with the genitourinary source (human papillomavirus and molluscum contagiosum virus), urinary excretion (BK and JC polyomavirus) or viruses possibly leaked into urine from bleeding and inflammation (Herpesvirus 6 and Anellovirus). As previously reported, excretion of polyomavirus is more commonly observed for JC than BK virus among nonimmunosuppressed individuals [28, 29], and excretion increases with immunosuppression [30]. Herpesvirus 6 is rarely excreted in urine [31]. During acute infection, some children with exanthema subitum may present sterile pyuria [32]. However, a likely source of significant number of viral reads in urine may be the sloughing off of cells in individuals with integrated copies of the HHV6 in the host genome – occurring in 0.5 to 1% of the population [33, 34].

### Gender

We observed differences in the microbiome content across sex (S4A Fig). The greatest differences (not significant after multiple testing correction) were greater numbers of sequence reads for *Lactobacillus* and *Prevotella* in women, and of *Enterococcus*, and *Pseudomonas* in men (S4B Fig).

## Discussion

This study provides a detailed view of the microbial metagenomes of urine specimens. The study departs from a classical analysis in that it maximizes a data-driven approach that extracts laboratory metadata features and matches them to metagenomic profiles. It provides an unbiased identification of the flora associated with samples colonized or contaminated with vaginal commensal organisms or local flora, and with samples associated with infection. Colonizing bacteria may be present at the urinary meatus, the distal urethra or along the entire urothelium. Where such bacteria reside cannot be determined from voided urine samples. Out study extends the identity of possible pathogens to include unconventional microorganisms and thus represents a new view of the nature of infection in the genitourinary region and an approach to the question of a normal urinary tract flora.

To evaluate those questions, we used both 16S rDNA and whole genome shotgun sequencing techniques. The former provided a greater sensitivity, as it identified bacterial species across the majority of samples and clinical groups. In contrast, less than half of the clean catch urine samples generated sequencing libraries for WGS. The basis for the lower sensitivity rests on the fact that WGS uses limited technical amplification of the nucleic acid content in the sample, thus more closely reflecting the proportionate masses contributed by microbes in the urinary metagenomes. WGS also provides a unique view of non-prokaryotic content of urine through the identification of eukarya - mainly Candida species, and of human viruses and phages. These differences notwithstanding, both sequencing techniques identify a substantial diversity of microbial species. WGS also provides a representation of virulence factors in the bacterial pool of the individual. Not unexpectedly, the analysis identifies differences in the microbial metagenome across sexes.

The study convincingly identifies high numbers of sequence reads of conventional uropathogens, but also proposes novel bacterial species associated with features of infection. It also challenges the cutoffs used to define infection: generally, 10^5^ colony forming units in culture. The quantitative nature of the WGS approach identifies traditional uropathogens at lower quantities in samples with features of infection. It identifies non-cultured/difficult to grow bacteria long discussed as a possible pathogenic organism, for example, *Alloscardovia* [35–38] and *Actinotignum* sp. [21]. WGS also provides a broader screen compared to the conventional urinary culture. For example, we identified sequence reads of *Ureaplasma* – a potential pathogen that requires dedicated culture systems or molecular testing. It is expected that the approach identify *Mycoplasma*, *Chlamydia* and other agents associated with sexually transmitted diseases.

Use of WGS also captures viral DNA sequence reads. The identification of viruses in the genitourinary tract is important because of the potential for transmission from local disease (e.g., HSV2, papillomaviruses), or because of the interest in monitoring of shedding (e.g., CMV, BK virus). WGS also identified shedding of common blood viruses such as Anellovirus (Torque teno virus) [39]. There is however limited information on the role of viruses as a cause of UTI [40]. Consistent with the work of Santiago-Rodriguez the al. [41], we observed the abundant presence of phages that match the presence of the cognate bacteria in urine. Metagenomic analyses could thus expand the understanding of the role of viruses as flora of the genitourinary tract.

The present study because uses specimens collected for clinical diagnostic purposes, but de-identified and considered medical waste. This limits in-depth understanding of the clinical setting beyond what can be established from the urine laboratory metadata. We propose that future studies on the urinary microbiome should use baseline unbiased microbial metagenome analysis to prospectively understand the nature of infection and of treatment response. We speculate that “Cluster 3” may to some extent include urine samples from individuals that received treatment with antibiotics. This cluster has the least amount of sequence reads in WGS, and the presence at low titers of classical uropathogens such as *Pseudomonas aeruginosa* or *Escherichia coli.* A systematic, prospective use of metagenomic tools may also shed light on the role of unknown and unconventional microorganisms in the urinary tract. Additional aspects that could be approached by urinary metagenomics are the characteristics of the urinary “normal flora” – as it is increasingly observed that the urinary tract may not be sterile. These studies could be performed via suprapubic collection of urine. Overall, the present study underscores that the current understanding of the etiology of UTIs can be improved through the combined used of unbiased clinical laboratory data and microbial metagenome analysis.

## Methods

### Study participants and urinalysis

A total of 121 human urine specimens were collected by the Pathology and Clinical Microbiology Laboratory of the Shady Grove Adventist Hospital (SGAH) in Rockville, Maryland. Details on the set of specimens and the urinalysis methods performed were described previously [5]. A total of 92 samples were collected from women, and 29 from men. The study was exempted from review by Internal Review Boards of the J. Craig Venter Institute (JCVI) and SGAH because the specimens were collected for diagnostic purposes and considered medical waste prior to use for the study. The urine samples left over after clinical urinalysis were de-identified prior to transfer to JCVI. Clinical laboratory records included gender and the results of urinalysis tests, such as presence of bacterial cells, red blood cells, leukocytes, epithelial cells and casts (assessed by phase contrast microscopy), nitrite concentration (associated with bacterial nitrate reduction) and leukocyte esterase activities (derived from the activity of white blood cell proteases and esterases released into urine).

### Sample processing

Urine specimens (5 to 30 ml) were stored at 4°C for up to 6 h after collection and centrifuged at 3,000 × g for 15 minutes at 10°C. Urinary pellets were washed twice with a 10-fold volume of PBS and frozen at −80°C until used for proteomic analyses as reported[5] or for microbiome and metagenomics analyses. On the day of DNA extraction, 300 μl of TES buffer (20mM Tris-Cl, pH 8.0, 2mM EDTA, and 1.2% Triton X-100) was added to a 5 to 25 μl urinary pellet sample. The sample was vortexed, incubated at 75°C for 10 min and cooled to room temperature. The suspension was supplemented with 60 μl chicken egg lysozyme (200 μg/ml), and 5 μl Linker RNase A, gently mixed and incubated for 60 min at 37°C. After addition of 100 μl 10% SDS and 42 μl Proteinase K (20 mg/ml), bacterial lysis was allowed to proceed overnight at 55°C. The DNA was extracted by adding an equal volume of phenol: chloroform: isoamylalcohol (25:24:1; pH 6.6), followed by vortexing and centrifuging at 13,000 RPM for 20 min. The aqueous phase was removed and transferred to a sterile microcentrifuge tube. The residual sample was then re-extracted by repeating the previous step. The aqueous phase was re-extracted with an equal volume of chloroform: isoamylalcohol (24:1) and centrifuged at 13,000 RPM for 15 min. The aqueous phase was transferred to a sterile microcentrifuge tube and 3 M sodium acetate (pH 5.2) was added at a 10% volume. The DNA was precipitated by adding an equal volume of ice cold isopropanol followed by incubation at −80°C for 30 minutes. Samples were then centrifuged at 16,100 x g for 10 min and the supernatant was removed. The pellet was washed with 80% ethanol and centrifuged again. After air drying, the DNA pellet was resuspended in Tris EDTA buffer in preparation for sequencing.

### Special phenotypic tests and proteomic methods

Previous work using the same samples focused on the integrated evaluation of urinalysis and proteomic data to diagnose UTI and inflammatory conditions in the genitourinary tract. The experimental shotgun proteomic methods were based on tryptic peptide analysis via nano-liquid chromatography tandem mass spectrometry (LC-MS/MS) with the high resolution high accuracy Q-Exactive mass spectrometer (V1.4, Thermo Electron) followed by computational searches of a database comprised of the combined sequences of the human proteome and 21 proteomes of microbial species known to colonize the human genitourinary tract [5]. Semi-quantitative proteomic data were obtained counting the peptide-spectral matches for a given proteins using the Proteome Discoverer™ software analysis tool (Thermo Electron) at a 1% peptide and protein false discovery rates. Quantitative analyses for the performance of phenotypic tests utilized the MaxQuant software tool [42]. The iBAQ protein values were computed for 35 proteins highly expressed in activated neutrophils, 32 proteins highly expressed in erythrocytes and five proteins highly expressed in squamous epithelial cells (cornifelin, cornulin, galactin-7, serpin B3, and mucin 5B) compared to the abundance of the urine-specific protein uromodulin. Summed iBAQ values then permitted the calculation of scores, which were termed the NAD (neutrophil activation and degranulation) score for neutrophil contents, the ERY (erythrocyte) score for red blood cell contents and the VCO (vaginal contamination) score for squamous epithelial contents [5]. These data allowed us to describe disease phenotypes for samples under investigation.

### Sequencing and analysis of 16S rDNA genes

DNA extracted from urine samples was amplified using primers that targeted the V1-V3 regions of the 16S rDNA gene [43]. These primers included the i5 and i7 adaptor sequences for Illumina MiSeq pyrosequencing and unique 8 bp indices incorporated into both primers such that each sample received its own unique barcode pair. The method of incorporating the adaptors and index sequences into primers at the PCR stage provided minimal loss of sequence data when compared to previous methods that would ligate adaptors to every amplicon after amplification. This method also allowed the generation of all sequence reads in the same 5’-3’ orientation. Using approximately 100 ng of extracted DNA, amplicons were generated with Platinum Taq polymerase (Life Technologies, CA) using the following cycling conditions: 95°C for 5 min for an initial denaturing step followed by 95°C for 30 sec, 55°C for 30 sec, 72°C for 30 sec for a total of 35 cycles followed by a final extension step of 72°C for 7 min then stored at 4°C. Once the PCR for each sample was completed, the amplicons were purified using the QIAquick PCR purification kit (Qiagen Valencia, CA), quantified fluorometrically using SYBR Gold Nucleic Acid Gel Stain (ThermoFisher Scientific), normalized, and pooled in preparation for bridge amplification followed by Illumina MiSeq sequencing using V3 chemistry dual index 2x300 bp format (Roche, Branford, CT) following the manufacturer’s protocol.

### Phylogenetic classification

16S rDNA amplicons were quality control using Infernal [44]. Only sequences identified as bacterial 16S using Infernal were considered for downstream steps. Bacterial 16S sequences were searched against SILVA (release 128) [45] using blastn [46] to initially determine the species found in the samples to include the corresponding SILVA reference sequences in a reference phylogenetic tree. Identified reference sequences were aligned using MAFFT [47] with the G-INS-i settings for global homology. A maximum likelihood reference tree was inferred under the general time-reversible model with gamma-distributed rate heterogeneity (GTR + Γ) using FastTree [48]. The 16S reads were mapped onto the reference tree using pplacer with the default settings. The number of sequences assigned to each node on the reference tree was normalized to the total number of sequences from the corresponding samples. The normalized abundances of the mapped reads were visualized using ggtree [49].

### Whole-genome shotgun sequencing

Nextera XT libraries were prepared manually following the manufacturer’s protocol (Illumina). Briefly, samples were normalized to 0.2 ng/μl DNA material per library using a Quant-iT picogreen assay system (Life Technologies) on an AF2200 plate reader (Eppendorf), then fragmented and tagged via tagmentation. Amplification was performed by Veriti 96 well PCR (Applied Biosystems) followed by AMPure XP bead cleanup (Beckman Coulter). Fragment size was measured using Labchip GX Touch high-sensitivity. For cluster generation and next generation sequencing, samples were normalized to 1 nM, pooled, and diluted to 8 pM. The paired-end cluster kit V4 was used and cluster generation was performed on an Illumina cBot, with pooled samples in all 8 lanes. Sequencing was performed on an Illumina HiSeq 2500 using SBS kit V4 chemistry. Median cluster densities (K mm2) were 908.5 for Nextera XT.

### Taxonomic assignments, microbial abundance, and virulence markers

Sequences were processed using the Human Longevity Inc. microbiome annotation pipeline as described in[50]. Briefly, after trimming adapter sequence, removing low quality bases, excluding reads shorter than 90 nucleotides, removing duplicated reads, reads were aligned to the human reference genome hg38 using BWA [51]. Reads that were mapped to hg38 were excluded from downstream analyses. Non-human reads were mapped to Human Longevity Inc. reference genomes database, which is composed of almost 19,023 NCBI reference genomes of bacteria, archaea, eukarya, and viruses. Successfully mapped reads were taxonomically classified using the Expectation Maximization algorithm [52]. The relative abundance of a reference genome was estimated as the genome coverage divided by the sum of all genome coverages. Non-human reads were assembled using IDBA-UD [53] and ORFs are predicted from assembled scaffolds with Metagene [54]. ORFs were compared against VFDB [23] to identify virulence factor genes with over 90% sequence identity.

### Dimensionality reduction of clinical data and clustering

The clinical laboratory metadata matrix was imputed for missing entries using MissForest [55]. Then principal component analysis (PCA) was conducted for a matrix of twenty clinical and sampling meta parameters (collection date, sex, urine appearance, urine volume, urine color, urine blood, hemoglobin presence with urine dipstick, red blood cells (RBCs), vascular injury score (ERY, see definition above), protein presence with urine dipstick, nitrate concentration, number of leukocytes, neutrophil activation and degranulation score (NAD, see definition above), complement system activity and coagulation, leukocytes microscopy, squamous epithelial cells [Epithelium], vagina contamination score [VCO, see definition above], urinary pellet appearance and color, urinary pellet volume and weight) from 121 individuals. The first two components from the PCA analysis, which explained 35% and 30% of the variance, were used to cluster the individuals using the partitioning around medoids (pam) method [56]. The optimal number of clusters was determined to be three using the silhouette method [57]. Microbial taxa were filtered for those with relative abundance ≥ 1e-4 in at least one individual.

### Data resources

The metagenomic sequence data will be deposited at NCBI under Bioprojects.

## Supporting information captions

**Figure S1.** Normalized abundance of 47 bacterial genera across the clinical groups using 16S rDNA.

**Figure S2.** Normalized abundance of 49 bacterial genera across the clinical groups using whole genome shotgun sequencing.

**Figure S3.** Correlation between 16S rDNA and whole genome shotgun sequencing. Each dot represents the median normalized abundance for 44 genera identified by the two sequencing approaches. Dots in reds represent those genus that exhibit greater than 2 standard deviation from the regression line, thus suggesting differential performance of one of the two techniques. The lack of correlation between *Raoultella* 16S rDNA and whole genome shotgun sequencing likely result from its recent reassignment as a separate genus from *Klebsiella*.

**Figure S4.** Differences across sex for selected microorganisms. A. Principal component analysis of female and male microbial metagenome showing differences along principal component PC1. B. Boxplot of genera that are nominally statistically different in number of read counts between female and male participants. These differences are not statistically different after multiple testing correction.

